# PhyloSOLID: Robust phylogeny reconstruction from single-cell data despite inherent error and sparsity

**DOI:** 10.64898/2026.02.04.703905

**Authors:** Qing Yang, Yuheng Liu, Jijian Yang, Xing Wu, Zhirui Yang, Yonghe Xia, Yunchao Zheng, Jinhong Lu, Mengdie Yao, Yiheng Du, Huan Liu, Nan Li, Yanmei Dou

**Author notes:** Correspondence (N.L.), (Y.Dou). These authors contributed equally (Qing Yang, Yuheng Liu).

## Abstract

While lineage tracing based on somatic mutations in single-cell sequencing data offers a powerful approach to reconstructing cellular histories in vivo, its reliability is fundamentally limited by pervasive technical artifacts—specifically, high error rates and data sparsity. These issues introduce false phylogenetic signals that corrupt tree topology and lead to spurious evolutionary conclusions. To overcome these limitations, we present PhyloSOLID, a phylogenetic algorithm designed to be inherently robust to these data imperfections. PhyloSOLID employs a progressive scaffolding strategy that begins with graph-based construction of a low-resolution, high-confidence backbone tree from reliable and uniformly covered mutations. This scaffold is then refined through the iterative integration of remaining data, guided by a Bayesian statistical model that penalizes phylogenetic inconsistencies to effectively separate the true evolutionary signal from technical artifacts. Benchmarking on both simulated datasets and multiple ground-truth datasets demonstrates that PhyloSOLID achieves superior lineage reconstruction accuracy over existing methods, for both single-cell RNA-seq and DNA-seq data. Additionally, a user-friendly web interface enables customized quality assessment, artifact removal, and interpretation of lineage structures. PhyloSOLID provides a powerful solution for decoding cellular evolution in developmental and disease contexts.

## Main

Somatic mutations serve as natural recorders of cellular lineage, providing a powerful means to reconstruct developmental and evolutionary trajectories in vivo^1–3^. The advent of single-cell sequencing has brought this promise into sharper focus by enabling the resolution of mutational histories at the cellular level^4,5^. However, the technical challenges inherent to these methods— including significant noise and extreme data sparsity—often obscure true biological signals. In single-cell whole-genome sequencing (scWGS), whole-genome amplification can introduce pervasive amplification artifacts^6^. In single-cell RNA sequencing (scRNA-seq), technical noise from library preparation and sequencing is compounded by biological processes such as RNA editing^7^. A universal challenge is allele dropout (ADO), where the signal from one allele is lost due to stochastic sampling, uneven coverage, or amplification bias. In scRNA-seq, this is further confounded by transcriptional dropout^8^, which occurs when a gene or an allele is not expressed in a cell, preventing the detection of its underlying genotype. These technical challenges lead to high error rates in somatic mutation calling and lineage tracing, especially when detecting mosaic mutations in non-cancerous cells, even with the most advanced detection pipelines^9^.

Despite the high error rates in somatic mutation detection, most existing phylogenetic tree construction methods for single-cell data incorporate these noisy sites without proper identification, filtering, or consideration during tree construction^10–13^. Additionally, most current methods fail to account for critical issues including allele dropout, extreme data sparsity, and cell-type-specific gene expression biases^14–17^. Consequently, the resulting phylogenetic trees are often inaccurate, distorting lineage relationships and complicating the interpretation of cellular evolutionary history (Fig. 1a-b). Given the importance of precise phylogeny reconstruction for understanding developmental processes and disease progression, there is a pressing need for error-tolerant methods that can effectively address the noise and technical artifacts inherent in single-cell sequencing data.

**Fig. 1:**
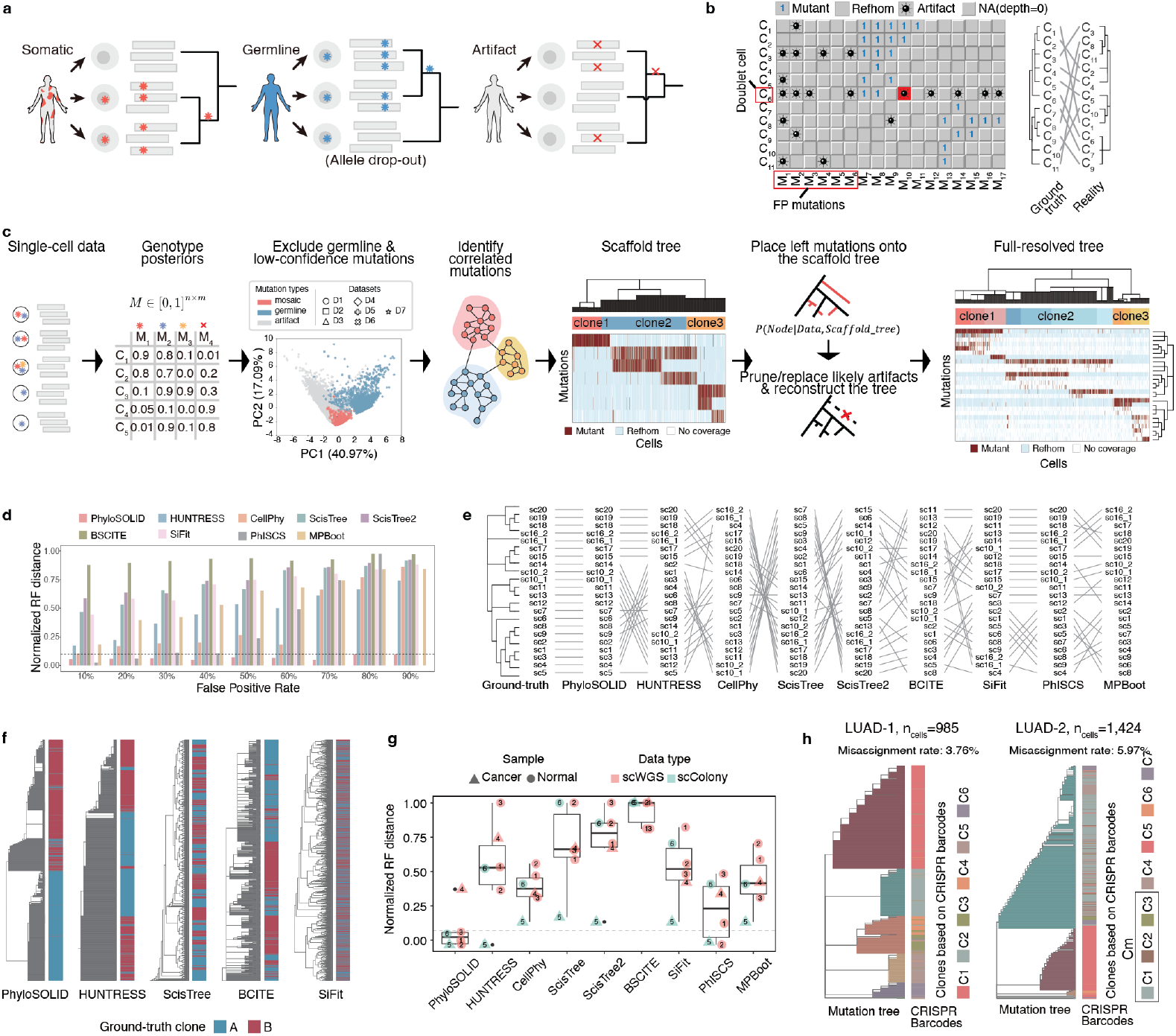
Development and benchmarking of PhyloSOLID. **a-b**, Single-cell phylogenetic tree structure can be distorted by several factors: germline variants misclassified as somatic mutations, erroneous mutation calls, allelic/depth dropout, and the presence of doublet cells. **c**, PhyloSOLID workflow. The PhyloSOLID framework takes a list of candidate somatic mutations and raw sequencing data as input. It then reconstructs an accurate phylogenetic tree by explicitly accounting for the major sources of error that distort tree structure. See Methods for details. **d**, Benchmarking phylogenetic inference with varying false-positive levels. PhyloSOLID and other tools were evaluated on a scDNA-seq dataset containing a controlled gradient of artifactual and germline variants (false positives). Phylogenetic accuracy was quantified using the normalized Robinson-Foulds (RF) distance. PhISCS execution failed at a 90% false positive rate. **e**, Tanglegram comparison at 50% false-positive rate. Compared with other tools, the tree inferred by PhyloSOLID remains highly congruent with the reference topology, as shown by the high node correspondence. **f**, Resolving distinct clones from sparse data. A sparse simulated cell-by-mutation matrix was generated, comprising two major clones (red and blue cells, rows). PhyloSOLID was benchmarked against other tools and correctly partitioned the cells into two independent clones. **g**, across six single-cell datasets with established ground-truth tree structures, PhyloSOLID consistently achieved superior phylogenetic accuracy. **h**, Accurate phylogenies from scRNA-seq data. Phylogenetic trees inferred by PhyloSOLID from somatic mutations in scRNA-seq data are highly congruent with the ground-truth phylogeny defined by CRISPR barcode mutations. In the LUAD-2 sample, no clone-specific mosaic mutations of C1-3 were detected. Instead, we identified a founding clone (Cm) that gave rise to the three descendent clones. “n_cells_” represents the number of cells.

To address this critical methodological gap, we developed PhyloSOLID, a robust phylogenetic reconstruction algorithm designed for single-cell somatic mutation data. Unlike conventional methods, PhyloSOLID incorporates several key innovations (Fig. 1c, Supplementary Fig. 1, Methods): (1) It learns the properties of artifact sites, germline variants, and true somatic mutations from multiple cross-validated, real-world data, excluding most germline and artifact sites prior to phylogeny construction (Supplementary Fig. 1, Supplementary Table 1)^18–23^. (2) It calculates genotyping posterior probabilities while accounting for allele imbalance issues. (3) It identifies correlated clones and mutations using a graph-based method, by explicitly modeling the inherent noise in the data (Supplementary Fig. 1b-e. (4) It first constructs a backbone tree of high-confidence clones supported by mutations from uniformly covered regions and then places remaining mutations onto this scaffold using a Bayesian penalty approach (Methods)^24^, preventing tree structure distortion from cell-type-specific gene expressions. (5) It refines the final topology by correcting misplaced variants, removing likely false-positive mutations and filtering out suspected doublet cells. (6) It provides an interactive website for user-customized quality control of the tree structure. Together, these features enable PhyloSOLID to overcome the inherent challenges of single-cell data, such as noise, sparsity and allele dropouts.

To evaluate performance under noise and data sparsity, we first used a scDNA-seq dataset^18^ (22 cells; 78 somatic mutations) and simulated varying noise levels by introducing artificial false-positive sites (Methods, Supplementary Table 2). PhyloSOLID consistently achieved superior accuracy, as measured by the normalized Robinson-Foulds (RF) distance^25^ to the ground-truth tree (Methods, Fig. 1d). Notably, even at a 90% false-positive rate, PhyloSOLID maintained a normalized RF distance below 0.1, with a representative example illustrated in Fig. 1e. To further test robustness under realistic data sparsity, we used a second simulated dataset comprising 50 mutations across 1,761 cells with a high dropout rate of ∼90% (Supplementary Table 3). Here, PhyloSOLID accurately reconstructed the ground-truth monoclonal phylogeny (Fig. 1f, Supplementary Fig. 2), whereas other widely used tools (e.g., HUNTREE, CellPhy, BCITE, ScisTree) erroneously inferred polyclonality.

After compiling a benchmark of eight ground truth datasets—comprising four scDNAseq benchmarks post whole-genome amplification^18,19^, two single-cell colony benchmarks with published phylogenies^20,21^, and two scRNAseq benchmarks with knock-in CRISPR barcode mutations^26^—we further evaluated the performance of PhyloSOLID against eight state-of-the-art methods (e.g., HUNTREE, CellPhy, ScisTree). Across all the scDNA-seq benchmarks, PhyloSOLID consistently and significantly outperformed other methods (Methods, Fig. 1f, Supplementary Fig. 3).

Notably, on two scRNA-seq benchmarks comprising over a thousand cells each, PhyloSOLID correctly assigned 94.03% and 96.24% of mutant cells to their original clones, as validated by the CRISPR barcode trees, achieving high phylogenetic accuracy despite the inherent noise and sparsity. In contrast, other tools exhibited high clone misassignment rates, ranging from ∼30% to 80% (Fig. 1g, Supplementary Fig. 4-6). Remarkably, using only somatic mutations, PhyloSOLID achieved phylogenetic resolution comparable to CRISPR-based barcoding across the scRNA-seq benchmarks tested. This result demonstrates that our method successfully unlocks the potential of somatic mutations for in vivo lineage tracing. Furthermore, the projection of cell lineages onto the gene expression UMAP revealed distinct transcriptional profiles for each clone (Supplementary Fig. 6), demonstrating that lineage history is coupled with changes in cell state and fate.

Phylogenetic trees reconstructed from single-cell data often involve large numbers of cells and mutations, along with additional layers of information such as genotype calls and cell-type annotations. These trees can be highly complex, making well-designed visualization schemes essential for meaningful interpretation. However, existing phylogenetic tools typically offer limited or inflexible visualization options—ill-suited to diverse scenarios such as trees with few mutations across many cells, or vice versa—and often require advanced coding skills for customization^27^. Furthermore, even robust methods like PhyloSOLID are not immune to limitations; residual false-positive mutations or doublet cells may still be incorporated and distort tree topology. Manual inspection and quality assessment are therefore crucial for accurate biological interpretation. Yet most current approaches deliver only a basic tree structure^15,28^, leaving users without adequate support for systematic quality evaluation.

To meet the demand for advanced visualization, we developed a suite of tailored tree visualization schemes (Fig. 2a–c). Each is designed to present multiple data layers—such as mutation number, cell-type annotations, and mutational profiles—in a clear, efficient, and publication-ready format. A rectangular tree layout is also provided to accommodate different user preferences (Fig. 2d). Beyond these general layouts, we developed specialized visualizations to suit different data structures and biological questions. These include trees that integrate single-cell mutation burdens, shared variants, and cell-type annotations (Fig. 2a); representations optimized for large numbers of mutations across few cells, with optional mutational signature overlay (Fig. 2b); and layouts tailored to large cell sets that emphasize a limited set of key driver mutations (Fig. 2c). Together, these options allow users to adapt tree visualization to their specific dataset and analytical goals.

**Fig. 2:**
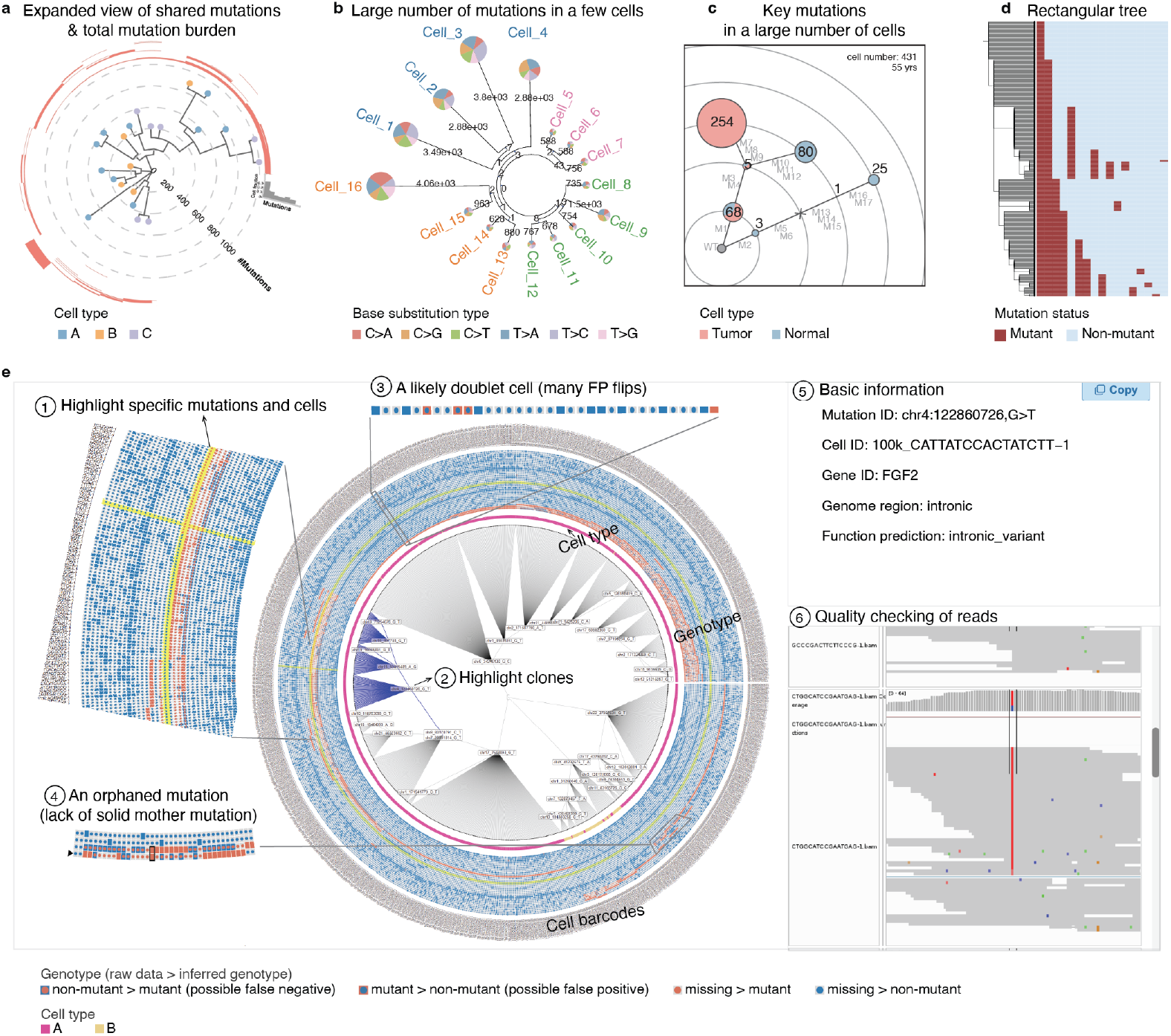
Advanced visualization and user-friendly interface. **a-c**, A suite of tailored tree layouts designed to intuitively integrate and visualize multiple layers of data—such as per-clade mutation burden, cell-type annotations, mutation profiles, and highlighting specific mutations or cells—in a clear and publication-ready format. **d**, A standard rectangular tree layout is also provided, offering flexibility to accommodate diverse user preferences. **e**, Interactive quality control and data exploration. A circular phylogenetic layout sorts and displays shared mutations across concentric rings, enabling direct visual comparison of input and output genotypes for each cell. This is integrated into an interactive web interface that allows users to perform point-and-click operations for in-depth analysis, such as inspecting read-level evidence, flagging potential false-positive mutations, and annotating doublet cells. An example of an “orphaned mutation” (black triangle) is illustrated here. This variant shares only one mutant cell with its assigned mother mutation yet causes many false-negative allele flips. This pattern suggests the lack of a true clonal mother mutation, likely resulting from a sequencing error, a doublet cell, or an independent mutation misassigned to this lineage.

For streamlined phylogeny quality control, we introduced a circular layout in which shared mutations are sorted and displayed across concentric rings, enabling direct comparison of input and output genotypes per cell (Fig. 2e). We provide an interactive web interface that enables users to perform a range of exploratory and quality-control operations. For example, users can visually inspect read-level evidence for mutations, flag potential false-positive sites, and mark questionable mutations and likely doublet cells—all through an intuitive, point-and-click environment. This interactive system greatly lowers the barrier to rigorous phylogenetic quality assessment, helping ensure robust interpretation of lineage relationships.

The PhyloSOLID source code is being prepared for public release. It will be available on GitHub (https://github.com/douymLab/PhyloSOLID) by March 1, 2026, or upon the manuscript’s formal acceptance, whichever comes first. An interactive website is accessible online at https://phylosolid.westlake.edu.cn.

## Discussion

The rapid expansion of single-cell sequencing has established somatic mutation and barcode-based lineage tracing as a central technique for decoding cellular fate, development, and disease. However, the intrinsic noisiness and sparsity of single-cell data—including pervasive allele dropout and low mutation-calling accuracy—pose a fundamental challenge to reliable phylogenetic inference. When unaddressed, these technical artifacts can be misinterpreted as biological signal, leading to incorrect reconstructions of cellular relationships and, consequently, erroneous biological conclusions. Here, we introduced PhyloSOLID, a computational framework designed to be inherently robust to these pervasive data imperfections.

The core strength of PhyloSOLID lies in its progressive scaffolding strategy, which prioritizes high-confidence phylogenetic signals before integrating more ambiguous data. By first constructing a robust backbone from reliable mutations and then iteratively refining it with a Bayesian model that penalizes phylogenetic inconsistencies, our method effectively disentangles true evolutionary history from technical noise. Extensive benchmarking on datasets with known ground truth demonstrated that PhyloSOLID consistently outperforms existing methods in reconstruction accuracy across both scRNA-seq and scDNA-seq modalities. Its performance on large-scale scRNA-seq datasets was particularly noteworthy, where phylogenetic trees built solely from somatic mutations recapitulated the clonal structure defined by independent CRISPR barcoding with remarkably high accuracy. This result underscores that somatic mutations, when analyzed with a sufficiently robust phylogenetic framework, can provide resolution for lineage tracing that rivals or even surpasses engineered barcoding approaches. Beyond constructing phylogenies from somatic mutations, PhyloSOLID is a flexible toolkit that can also leverage barcode mutations for phylogenetic inference.

Looking forward, the need for reliable single cell phylogenetics will become ever more pressing. PhyloSOLID meets this need by providing a statistically rigorous, error-aware framework for lineage reconstruction. To maximize its accessibility and utility for the broader research community, we have complemented the core algorithm with a user-friendly web interface for quality assessment, artifact removal, and lineage interpretation. We anticipate that PhyloSOLID will become an essential tool, empowering researchers to extract trustworthy evolutionary insights from the complex and noisy landscape of single-cell data and thereby ensuring that the burgeoning field of cellular lineage tracing is built upon a foundation of computational rigor.

## Data availability

All scWGS and scRNA-seq data used in this study are publicly available under the following accession codes: dbGaP: phs001485.v3.p1, SRA: SRA053195, NDA: NDAR#2330, EGA: EGAD00001007032, and GEO: GSE144239, GSE234814 and GSE161363.

## Code availability

PhyloSOLID is implemented in Python and R and is licensed under the MIT License. The source code, documentation and examples are available on GitHub at https://github.com/douymLab/PhyloSOLID. The PhyloSOLID source code is being prepared for public release. It will be available on GitHub by March 1, 2026, or upon the manuscript’s formal acceptance, whichever comes first.

## Acknowledgements

This work was supported by the Westlake Laboratory of Life Sciences and Biomedicine (Hangzhou 310024, Zhejiang, China), under the grant “Key R*j*D Program of Zhejiang” (2024SSYS0032) as well as the National Natural Science Foundation of China (32270682) to Y.D. We thank the High-Performance Computing Center for technical support. We gratefully acknowledge the creators and submitters of the datasets used in this study. Data used in this study under the dbGaP accession phs001485.v3.p1 was generated by Drs.

Christopher A. Walsh and Peter J. Park with funding from NINDS grant R01NS032457. The public datasets were obtained from dbGaP at http://www.ncbi.nlm.nih.gov/gap, EGA at https://ega-archive.org/, NDA at https://nda.nih.gov/ and GEO at https://www.ncbi.nlm.nih.gov/geo/.

## Author contribution

Y.Dou conceived and supervised the project and secured the funding. N.L. supervised the website construction. Q.Y. and Y.L. both made significant contributions to the project, with Q.Y. developed the software and accomplished bioinformatics analysis, and Y.L. constructed the website with help from J. Y. X.W., Y.X, Z.Y., J.L, Y.Z., J.L., M.Y., Y.Du and H.L. assisted in single cell genotyping, variant detection, and Y. X. assisted in visualization of phylogenies. Y.Dou and Q.Y. wrote the manuscript. All authors carefully reviewed and approved the final manuscript.

## Inclusion *j* Ethics statement

This study was conducted in accordance with ethical guidelines and principles. It did not involve human participant or animal subjects. There are no sequencing data generated in this study. The authors ensure that the study adheres to the highest ethical standards in research, data generation, and usage.

## Methods

### The PhyloSOLID framework

#### 1. Pre-phylogeny genotyping and variant filtering

We collected eight single-cell benchmarks, comprising six scDNA-seq benchmarks with orthogonally validated ground-truth mutations (Supplementary Table 4) and two scRNA-seq benchmarks with ground-truth CRISPR barcoding trees (Supplementary Table 5-6), from public sources. Mutations were genotyped using MosaicSC^24^, a Bayesian approach that our lab have developed, accounting for single-cell-specific challenges such as allelic imbalance, sequencing errors, and population allele frequencies. Following genotyping, we applied a logistic regression classifier trained with an “leave-one-donor-out” approach to filter artifacts and germline variants. This model distinguished true mosaic mutations based on 10 (for scDNA-seq) or 14 (for scRNA-seq) features derived from raw sequencing reads, including mismatch frequency, base quality, read alignment position, etc. (Supplementary Table 1). We further excluded mutations with a low mutant-allele fraction (VAF < 1%) or a population allele frequency >0.1% in gnomAD. For scRNA-seq-derived mutations, we further excluded those mapping to known sites in three classical RNA editing databases^29–31^ or residing within allele-specific expressed (ASE) genes^32^, those with significantly imbalanced base-substitution profiles, and those with low UMI consistency. Of note, when BAM files are unavailable, PhyloSOLID offers a practical alternative by constructing phylogenies directly from a cell-by-mutation matrix.

#### 2. Leveraging clonal architecture to further filter germline variants

To distinguish true somatic mutations from germline variants with allele dropout in single-cell DNA sequencing data, we leverage the fundamental principle that genuine somatic mutations exhibit structured patterns of co-occurrence and mutual exclusivity reflective of underlying clonal architecture. Specifically, mutations belonging to the same subclone tend to co-occur within the same group of cells, while those originating from distinct, mutually exclusive subclones are distributed across different cell populations. In contrast, germline variants affected by stochastic allele dropout demonstrate no systematic association with clonal groups; instead, they appear randomly distributed across cells and show no consistent correlation with other mutations, resulting in a nonspecific and biologically incoherent pattern.

#### 2.1) Identification of mutations with co-occurrence patterns

Let *n* and *m* be the number of cells and mutations retained after initial filtering, respectively. The final matrix used for analysis is *I* = {0, 1, *NA*}^*n*×*m*^, where *NA* values are excluded from calculations. This matrix is derived from three input matrices:

1. *P*_*i,j*_ ∈ [0,1]^*n*×*m*^: the matrix of somatic posterior probabilities, where *P*_*i,j*_ represents the probability that the somatic mutation *j* is present in cell *i*. Somatic mutation likelihoods were calculated based on a beta-binomial sampling model.
2. *M*_*i,j*_ ∈ [0,1]^*n*×*m*^: the mutant allele frequency matrix, where *M*_*i,j*_ represents the mutant allele frequency of somatic mutation *j* in cell *i*.
3. 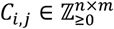 : the sequencing coverage matrix, where *C*_*i,j*_ represents the number of reads (coverage depth) at the locus of mutation *j* in cell *i*.

For each mutation *j* in each cell *i*, an entry in the binary matrix is defined as:

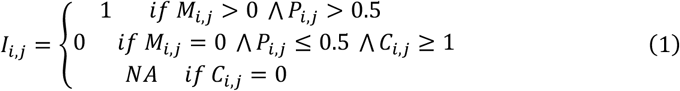

For each mutation *j*, the set of cells in which it was detected was defined as:

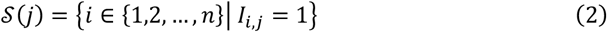

Mutation processing order: To ensure robust phylogenetic reconstruction, mutations are processed in descending order of their cellular prevalence |𝒮(*j*)|, with higher-confidence mutations processed first. This sorting strategy prioritizes mutations with stronger phylogenetic signals for initial tree construction.

For any pair of mutations *j*_1_ and *j*_2_, the following contingency counts were computed:

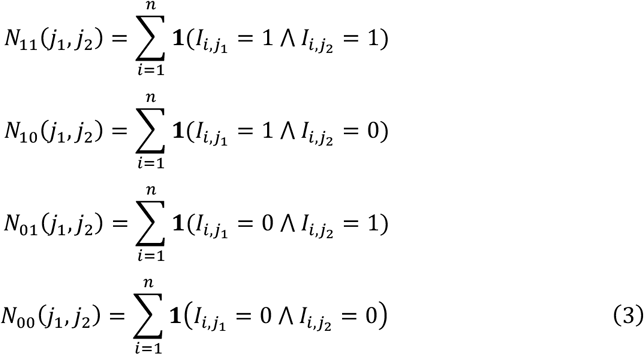

where **1**(·) was the indicator function, returning 1 if its argument is true and 0 otherwise.

We applied the Jaccard index to measure the similarity between the sets of cells containing each mutation:

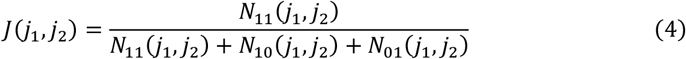

Jaccard index was selected for scRNAseq data because this metric ignores double-negative cells (*N*_00_), which is suitable for sparse scRNAseq data. However, this measure alone is insufficient for identifying parent-child clonal relationships, as the Jaccard index between a parent and its subclone may not necessarily be high. To specifically capture such hierarchical relationships, we introduced an additional metric defined as follows:

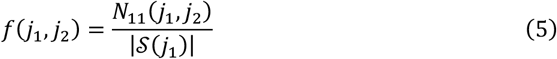

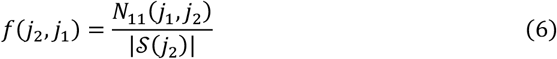

The Two mutations above, *j*_1_ and *j*_2_, were considered correlated if they satisfied either of the following criteria:

1. *N*_11_(*j*_1_, *j*_2_) > 0 ∧ *J*(*j*_1_, *j*_2_) ≥ 0.08,
2. *N*_11_(*j*_1_, *j*_2_) > 0 ∧ 0 < *J*(*j*_1_, *j*_2_) < 0.08 ∧ max (*f*(*j*_1_, *j*_2_), *f*(*j*_2_, *j*_1_)) ≥ 0.5.

#### 2.2) Exclusion of likely germline variants

To systematically identify and exclude germline variants misclassified as somatic mutations, we employed an exhaustive search approach that evaluates the patterns of co-occurrence and mutual exclusivity across all mutations.

A “founder mutation” is defined as a genetic alteration that initiates a clonal expansion and is subsequently present in all cells of that clone and all its subclones. To systematically identify such mutations and their associated clonal groups, we proceed as follows:

For each mutation *r* ∈ {1, …, *m*} with a mutant cell fraction (MCF) >5%, we consider it as a candidate founder mutation. For each such candidate *r*, we identify the set of mutations 𝒥, that are significantly correlated with *r*, as described in Section **2.1**.

The underlying rationale is that if *r* is a true founder mutation, the set 𝒥, likely represents subsequent mutations that occurred within the clone founded by *r*, and thus should exhibit a strong pattern of co-occurrence with *r*.

##### 2.2.1) Genotype imputation of the “founder mutation”

In theory, these correlated clones descend from a single founder clone. Consequently, all daughter clones are expected to inherit the founder’s mutations. Assuming *r* as the founder mutation of these correlated clones, we created a combined mutation set that includes both *r* and all mutations significantly correlated with it:

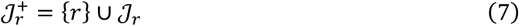

For each cell, we then quantified *q*_*i*_, the number of mutations from the combined set 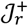 that are present in that cell:

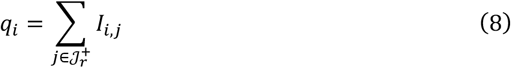

Where:

- *q*_*i*_: The number of mutations from the set 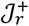 that are present in cell *i*. Impute founder status: A cell is classified as carrying the founder mutation *r* if it possesses at least two mutations from the set 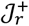. This threshold helps mitigate false negatives caused by allele dropout or sequencing errors affecting a single mutation.

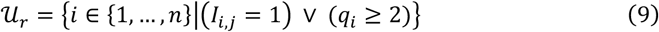

Where:
- 𝒰,: The set of cells inferred to belong to the clone founded by mutation *r*. Let 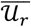 denote the complement of 𝒰, with respect to covered cells for any mutation *j*, i.e.,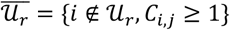. For each mutation 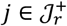, we calculated the total number of mutant cells outside the inferred founder clone:

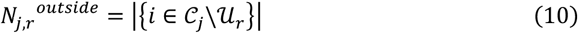

Where:
- 𝒞_j_ denotes the set of *j*-mutant cells.

Since all cells carrying mutation 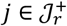 are descendants of the founder clone, no mutant cells should exist outside of it. We then defined a false-positive penalty score for mutation *j* by normalizing 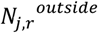 by the total number of *j*-mutant cells:

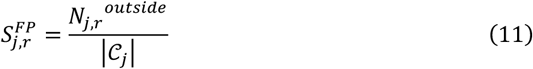

If r and *j* are both real mosaic mutations, then 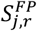 will be low. Conversely, a high score suggests otherwise.

##### 2.2.2) Enumeration over all candidate founder mutations and identification of likely germline variations

The above procedure (Sections 2.2.1 and 2.2.2) was repeated for each mutation *r* ∈ {1, …, *m*}. For each mutation r, we calculate a summary leakage score: (cv\std\mean, fig)

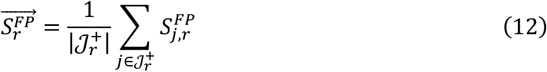

This score represents the average phylogenetic inconsistency of all mutations in the clonal set 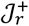 founder by *r*.

To identify germline variants, all candidate mutations *r* are sorted in descending order of 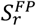. Germline variants form a distinct group at the top of this sorted list. We identify the cutoff point by calculating differences between consecutive sorted scores:

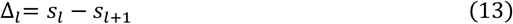

where *s*_*l*_ represents the *l*-th highest score in the sorted sequence. Mutations preceding a large drop in scores (Δ_*l*_> *μ*_Δ_ + 2*σ*_Δ_, where *μ*_Δ_ and *σ*_Δ_ are the mean and standard deviation of all differences) are classified as germline.

These identified germline variants are designated as ℳ_*het*_, placed at the root of the clonal tree, and excluded from subsequent lineage reconstruction. This approach effectively separated germline variants from real mosaic mutations.

### 3. Scaffold tree construction using uniformly covered mutations

Following the exclusion of germline mutations, we proceed to identify high-confidence somatic mutations located in genomic regions with ubiquitous expression and uniform coverage. These mutations serve as backbone features for initial phylogenetic reconstruction.

#### 3.1) Initial filtration based on genotype posterior and mutant reads count

We begin with the matrices defined in Section 2.1 and apply additional quality filters to ensure reliable detection of ubiquitously expressed mutations.

Cell Filtering:

The set of retained cells is defined as those containing sufficient mutation evidence:

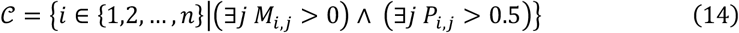

where *M* and *P* are the mutant allele frequency and somatic posterior probability matrices. Mutation filtration:

For each mutation *j*, we compute its average mutant allele frequency (MAF) across cells where it is detected. The set of cells where mutation *j* is detected is defined as:

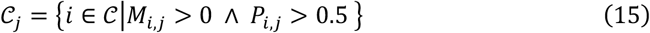

The average MAF for mutation *j* is:

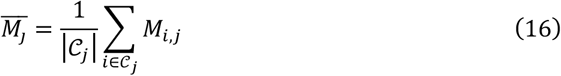

The set of retained mutations is:

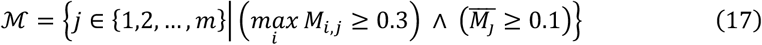

The filtered matrices are then:

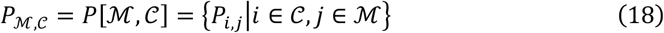

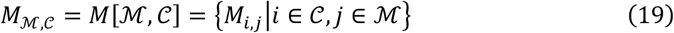

#### 3.2) Selection of uniformly covered mutations

To identify high-confidence mutations within ubiquitously expressed genomic regions, we applied two complementary coverage-based filters.

For mutation *j* in cell type *t*, the proportion of cells lacking sequencing coverage is denoted as:

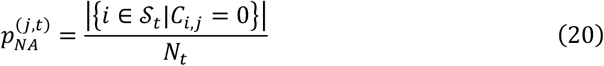

Where:

- 𝒮_*t*_ denotes the set of all cells that belong to cell type *t*;
- *C*_*i,j*_ denotes the sequencing coverage for mutation *j* in cell *i*;
- *i* denotes a cell *i*;
- *N*_2_ denotes the total number of cells of cell type *t*.

For each mutation *j*, we define *N*_*j*_ as the number of cell types in which a sufficient proportion of cells exhibit sequencing coverage:

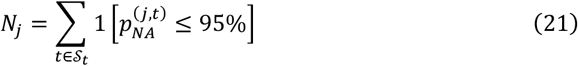

Where:

- 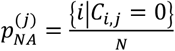 NA proportion for mutation *j* across all cells.
- 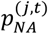 NA proportion for mutation *j* in cell type *t*;

Based on measurements from real-world single cell sequencing data, mutations are retained for subsequent analysis as candidate scaffold mutations if they meet:

- Present in >10% cells across all cells when there’s only one single dominant cell type, or no cell type annotation information is available.
- Present in ≥2 cell types (*N*_*j*_ ≥ 2).

For each mutation *j*, we first required that the median sequencing depth across all cells is at least 1x.

We then calculated the coefficient of variation (CV) to assess coverage uniformity. For mutation *j* in cell *i*, we defined a binarized coverage indicator:

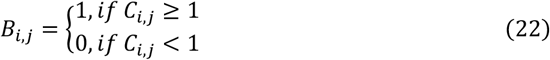

Where:

- *C*_*i,j*_ indicates the read coverage in cell *i* at the genome location of mutation *j*;

*B*_*i,j*_ is the binary indicator of whether mutation *j* in cell *i* has sufficient read coverage.

For each mutation *j*, the mean read coverage along with the standard deviation, and coefficient of variation are computed as follows:

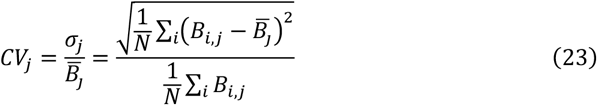

Where:

- *C*_*i,j*_ denotes the sequencing coverage for mutation *j* in cell *i*;
- *N* is the total number of cells.
- 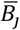 is the mean of *B*_*i,j*_ across all cells for mutation *j*;
- σ_*j*_ is the standard deviation of *B*_*i,j*_ across all cells for mutation *j*;
- *CV*_*j*_ is the coefficient of variation for the coverage indicator of mutation *j*.

Based on the measurements in real-world data, only mutations with *CV*_*j*_ < 6 were retained as candidate scaffold mutations.

#### 3.3) Noise-robust quantification of pairwise mutation correlations

Theoretically, two somatic mutations in an individual can only have one of four phylogenetic relationships: A is ancestral to B, B is ancestral to A, they co-occur on the same node, or they arise on independent branches. Consequently, if two mutations share an ancestral relationship or occur on the same node, they are supposed to co-exist in at least a subset of cells. In contrast, mutations on independent branches are not expected to to-exist in any single cell. Therefore, the co-occurrence patterns of mutations across single cells can be used to identify correlated mutations.

However, single cell sequencing data are inherently noisy. Sequencing errors and the presence of doublet cells can obscure the true phylogenetic relationships between mutations. To establish robust co-occurrence patterns while mitigating stochastic effects, we developed a consensus approach based on ensemble clone identification through randomized runs.

Beginning from each scaffold mutation *j* ∈ *M*_*S*_, we sequentially evaluated its correlation with every other scaffold mutation using the method described in Section 2.1. Mutations found to be correlated with *j* were assigned to the same clonal population with *j*. This process was repeated iteratively until all mutations were assigned to some clone. To ensure robustness, the scaffold mutation order was randomly shuffled, and the entire procedure was repeated 100 times.

The occurrence frequency of each distinct clone *C* across 100 runs were then calculated:

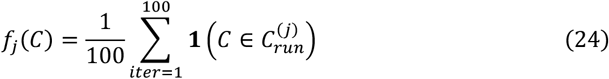

Only clones with *f*_*j*_(*C*) > 0.1 were retained, denote as 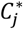. The weight for each clone *C* to exist was then computed as:

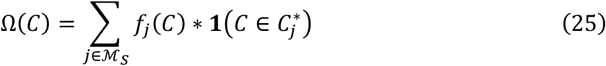

For each clone with |*C*| > 2, we extracted all 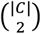 unordered mutations pairs. Each pair (*j, j* ^′^) inherited weight Ω(*C*) from its parent clone. The consensus weight for each distinct mutation pair was obtained by summing contributions from all clones containing both mutations:

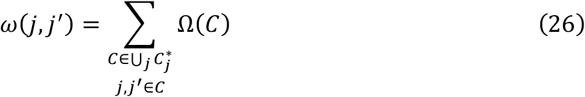

For singleton clones where |*C*| = 1, although they do not contribute to edge weights, their weights Ω(*C*) are retained and provide valuable information about mutation independence. Mutations with high singleton weights typically exhibit weak correlations with other mutations and are more likely to form distinct clonal groups in subsequent clustering analysis.

We constructed a weighted graph *G*_*C*o*nsensus*_ = (𝒱, ℰ, 𝒲) where:

- 𝒱 = ℳ_*S*_: vertices represent mutations.
- ℰ = {(*j, j*^′^)|*ω*(*j, j*^′^) > 0}: edges connect co-occurring mutations.
- 𝒲(*e*) = *ω*(*j, j*^′^): edge weights reflect co-occurrence strength.

This graph serves as the input for subsequent community detection. In this way, the relationship between each mutation pair was robustly quantified, making the approach resistant to stochastic noise.

#### 3.4) Partitioning of scaffold mutations into clonal groups via weighted graph clustering

The weighted consensus correlation graph *G*_*Consensus*_ was partitioned into maximal mutation groups using the Leiden community detection algorithm. This approach identifies phylogenetically coherent groups of mutations that exhibit strong internal correlations and minimal external correlations, with each group defining the mutational signature of a distinct clonal population.

The algorithm produces a complete partition of the scaffold mutation set 𝒢 = {*g*_1_, *g*_2_, …, *g*_K_}, where 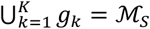 and *g*_*k*_ ∩ *g*_*k*′_ = ∅ for *k* ≠ *k*^′^. Each mutation group *g*_*k*_ defines the characteristic mutational profile of a putative cellular subpopulation, representing clones that likely evolved through distinct evolutionary trajectories.

From each phylogenetically coherent mutation group *g*_*k*_ ∈ 𝒢, we selected a single backbone mutation to serve as the phylogenetic anchor and operational founder of that clonal group.

The founder mutation of a clone is expected to be an early event, present in the vast majority of cells within that clonal lineage. Therefore, for each group *g*_*k*_, we selected the mutation with the highest Mutant Cell Fraction (MCF), defined as the fraction of cells with adequate sequencing coverage that harbor the mutation:

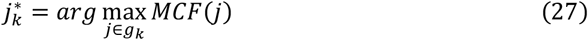

where 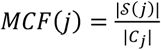. Here, 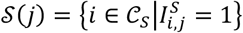 is the set of scaffold cells supporting mutation *j*, and |*C*_*j*_| denotes the total number of scaffold cells with non-zero sequencing coverage at locus *j*.

The stringent, coverage-based filtration applied to define the scaffold set ℳ_*S*_ (Section 3.3) ensures that all candidate mutations have low genome-wide missing data rates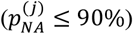 or are robustly detected across multiple cell types (*N*_*j*_ ≥ 2). This pre-processing step mitigates the risk of selecting mutations with spuzriously high MCF due to technical artifacts, thereby guaranteeing that a high MCF is a reliable indicator of high cellular prevalence.

The final set of backbone mutations is defined as the union of all group representatives:

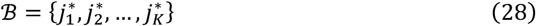

These backbone mutations provide a robust set of phylogenetically informative markers for the initial scaffold tree construction. This selection provides the foundational framework for the phylogenetic tree. Notably, the designation of an initial founder mutation is operational for the initial tree building and may be subject to re-evaluation during subsequent steps when placing additional mutations, allowing for the integration of more complex phylogenetic signals.

Following the selection of backbone mutations as phylogenetic anchors, we define the corresponding backbone clones that constitute the fundamental phylogenetic units in our reconstruction framework.

Definition 3.3.4.1 (Backbone Clone). For each backbone mutation 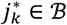 representing group *g*_*k*_, the corresponding backbone clone 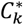 is defined as the set of scaffold cells exhibiting the characteristic mutation pattern of that phylogenetic lineage:

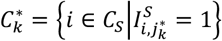

To robustly assign cells to backbone clones while accounting for missing data and technical dropouts, we employ a group-wise imputation strategy. For each backbone mutation 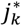, an imputed clone vector *V*_*k*_ is constructed by integrating phylogenetic signals from all mutations within group *g*_*k*_:

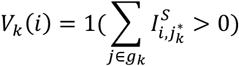

where 1(·) is the indicator function. This approach enhances detection sensitivity by leveraging phylogenetic coherence within each group.

Cells may exhibit mutation patterns consistent with multiple backbone clones due to phylogenetic relationships or technical artifacts. To resolve such ambiguities, a maximum-support assignment strategy is applied. Let 𝒜(*i*) = {*k*|*V*_*k*_(*i*) = 1} denote the set of candidate backbone clones for cell *i*. The final assignment is determined by:

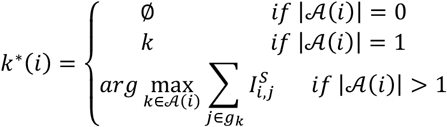

In case of ties, the backbone mutation with the highest expression level in the cell is selected:

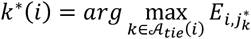

where *E*_*i,j*_ denotes the expression level of mutation *j* in cell *i*, and 𝒜_*tie*_(*i*) is the set of clones with equal maximal support.

The backbone clone matrix *B*_*Cl*0*ne*_ is then constructed as:

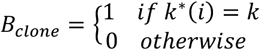

This matrix provides a robust binary representation of clonal membership, forming the basis for subsequent phylogenetic tree construction. The assignment procedure ensures that each cell is assigned to at most one backbone clone, leveraging phylogenetic coherence to enhance robustness against technical noise while resolving ambiguities using biologically meaningful criteria.

### 4. A Bayesian penalty approach to place left mutations onto the scaffold tree

The scaffold tree reconstruction proceeds iteratively, starting from the backbone tree *T*_*B*_ containing only the backbone mutations ℬ. Each non-backbone mutation *a* ∈ ℳ_*S*_ \ ℬ is sequentially integrated into the current tree *T*, which evolves during the reconstruction process. For each candidate mutation *a*, we evaluate its phylogenetic relationship to every possible placement position within *T* by computing a binary discordance penalty based on co-occurrence patterns and a comprehensive Bayesian penalty that accounts for phylogenetic structure and uncertainty.

#### 4.1) Identification of candidate nodes to place the mutation

For a given non-backbone mutation *a*, we comprehensively evaluate all phylogenetically permissible placement positions within the current tree *T*. Candidate positions *b* ∈ *T* includes: (i) any existing node in *T*; (ii) a new node introduced along any existing branch, thereby splitting it into two segments; (iii) a new parent node unifying two or more descendants from an existing node; or (iv) a new leaf node attached to any existing node. This exhaustive consideration ensures that all potential evolutionary relationships—including those requiring expansion of the tree topology—are evaluated.

To systematically manage these placement options, we employ an anchor-based approach. For each candidate node *b* ∈ *T* (which represents a distinct branch point or phylogenetic position within the tree) and non-backbone mutation *a* ∈ ℳ_*S*_ \ ℬ, we compute the standard contingency counts *N*_11_(*b, a*), *N*_10_(*b, a*), *N*_01_(*b, a*) and *N*_00_(*b, a*) as defined in Equation 3. To account for uncertainty in mutation calls derived from posterior probabilities *P*_*i,j*_, we define for each cell in these contingency categories a minimum and maximum possible penalty contribution, based on whether the observed posterior is close to the 0.5 threshold or confidently assigned.

The total binary discordance penalty for the pair (*a, b*) is defined as:

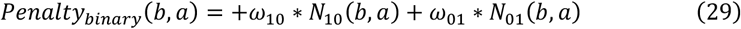

where the weights are assigned as follows:

- *ω*_10_ = 1: Full penalty for each cell in *N*_10_, representing a potential false positive (mutation *a* observed but one expected under position *b*;
- *ω*_01_ = 0.1: A reduced penalty for each cell in *N*_01_, representing a potential false negative (mutation *a* not observed but expected under position *b*), reflecting the higher empirical rate of false negatives in single-cell data.

For each contingency count, we compute the minimum and maximum possible penalty per cell, considering the uncertainty in posterior probability values around the 0.5 threshold:

- For a cell in *N*_10_(*b, a*): the penalty contribution ranges from 0.5 (if *P*_*i,j*_ is just above 0.5) to 1 (if *P*_*i,j*_ = 1).
- For a cell in *N*_01_(*b, a*): the penalty contribution ranges from 0.05 (if *P*_*i,j*_ is just below 0.5) to 0.1 (if *P*_*i,j*_ = 0).

Thus, the overall minimum and maximum penalties for the pair (*b, a*) are:

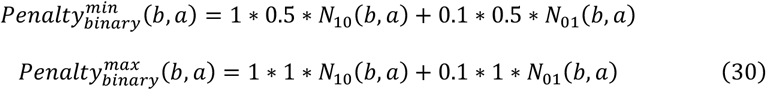

To identify candidate anchors, we compare the penalty intervals across all possible *b* ∈ *T*. We first identify the mutation *b*^∗^ with the smallest minimum penalty:

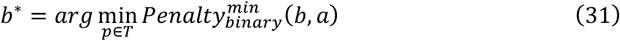

The candidate set is then constructed by including all positions *b* for which the minimum penalty is no greater than the maximum penalty of this optimal candidate:

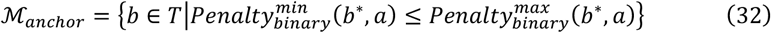

This approach ensures that we consider not only the position with the absolute smallest penalty, but also any other positions whose minimum penalty falls within the uncertainty range of the best candidate.

For each candidate anchor *b* ∈ ℳ _*anchor*_, we consider two primary phylogenetic hypotheses:

- Placement on position *b* : implying an identical phylogenetic origin (*a* = *b*), suggesting that observed differences (*N*_10_, *N*_01_) are due to sequencing errors.
- Placement as a direct descendant of position *b* : implying a parent-child relationship (*a* ⊂ *b*), where *a* is a subclone of *b*.

Additionally, placements that introduce new nodes along the branch leading to *b*, or that unify *b* with other siblings under a new parent node, are evaluated through an extension of this anchor-based framework, leveraging the same penalty structure to ensure phylogenetic consistency.

#### 4.2) A Bayesian approach to penalization

We applied a Bayesian approach to place new mutations onto the scaffold phylogenetic tree. The optimal placement was identified by maximizing the posterior probability of the mutation’s location, conditional on the scaffold tree structure and the observed data. This is formally equivalent to minimizing a penalty function defined as the negative log-posterior probability. The Bayesian function is as follows:

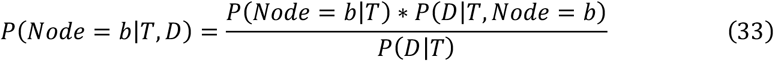

Where:

- *Node* : a candidate phylogenetic position in tree *T* where mutation might be placed.
- *D* = {*P*_1,*a*_, *P*_2,*a*_, …, *P*_*n,a*_}: the somatic posterior probabilities vector for mutation across all cells.
- *P*_*i,a*_: probability that mutation *a* is present in cell *i*.

Taking the negative logarithm yields the penalty function:

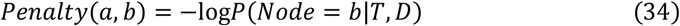

Expanding this expression:

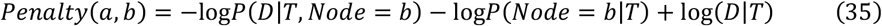

Since *P*(*D*|*T*) is constant across candidate placements for the current tree:

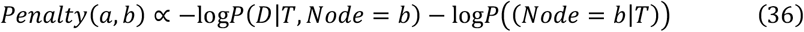

Assuming a uniform prior over *N*_*nodes*_ possible placement nodes in the current tree *T*:

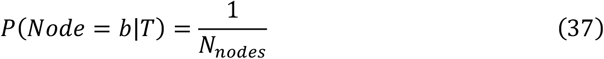

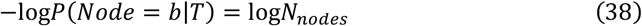

The data likelihood factors over cells based on the current tree structure:

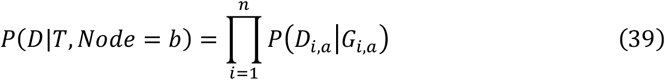

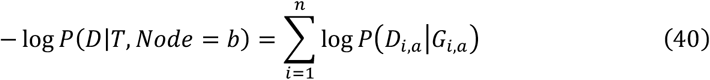

Where:

- *G*_*i,a*_: expected genotype (0 or 1) given placement at *Nodes*;
- *D*_*i,a*_: observed data (posterior probability *P*_*i,a*_).

#### 4.3) Mutation placement by minimizing the penalty function

The theoretical framework translates to the practical penalty function:

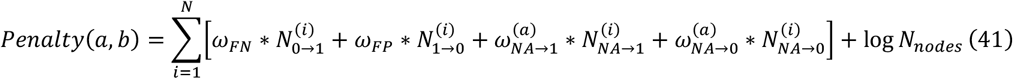

Binary discordance indicators (0 or 1):

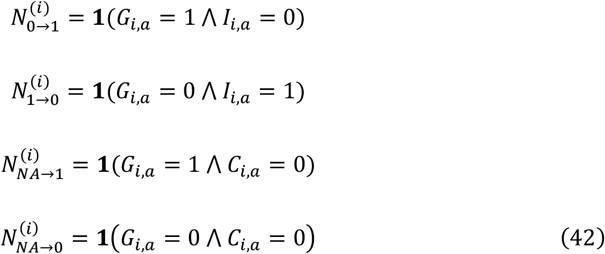

Where:

- Each indicator function **1**(·) returns 1 if the specified condition is true, 0 otherwise.
- 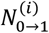: indicates if cell *i* has a false negative (expected mutant but observed reference).
- 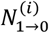 : indicates if cell *i* has a false positive (expected reference but observed mutant).
- 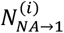 : indicates if cell *i* has missing data imputed as mutant.
- 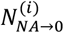 : indicates if cell *i* has missing data imputed as reference.

Weight computation from posterior matrix *P*:

- *ω*_*FP*_ = − log(*P*_*i, a*_): penalty for false positives.
- *ω*_*FN*_ = 0.1 ∗ (− log(1 − *P*_*i, a*_)): penalty for false negatives, scaled by empirical error ratio.
- 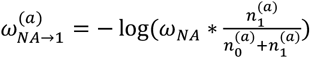: penalty for imputing NA as mutant.
- 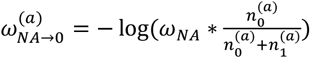: penalty for imputing NA as reference.

Empirical error ratio incorporation:

Based on extensive analysis of single-cell phylogenetic trees from real datasets, we observed that false negative errors occur approximately 10 times more frequently than false positive errors. This empirical observation (*fnfp*_*ratio* ≈ 0.1) is incorporated by scaling the false negative penalty, ensuring the penalty function reflects actual error patterns in single cell sequencing data. Both penalties maintain their log-probability interpretation while accounting for real-world error distributions.

Where:

- 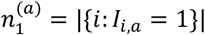: number of cells with observed mutant allele.
- 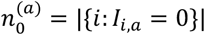: number of cells with observed reference allele.
- *ω*_*NA*_ = 0.001: global dropout weight for typical datasets.

Additionally, we incorporate a complexity penalty based on the Bayesian Information Criterion (BIC) to avoid overfitting:

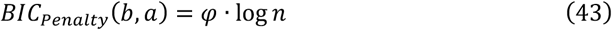

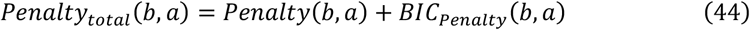

Where:

- *φ*: increase in number of parameters when adding *a*;
- *λ*: regularization parameter for complexity control.

The mutation *a* is placed at the candidate node *b*^∗^ that minimizes the total penalty:

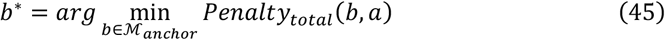

If ℳ _*anchor*_ = ∅, the mutation is placed at the root of the tree, subject to a root-specific penalty calculation.

After processing all non-backbone mutations through this iterative placement procedure, the resulting tree structure represents the complete scaffold tree *T*_G_, containing all scaffold mutations at their phylogenetically consistent positions.

The scaffold tree *T*_G_ provides a robust evolutionary framework grounded in high-confidence mutations from ubiquitously expressed regions, serving as the foundation for subsequent phylogenetic analysis and placement of additional mutations.

### 5. Prune *j* replace likely artifacts and reconstruct the tree

Following initial tree construction, each mutation was evaluated using five quality metrics to identify potential misplacements:

a. The false-positive flip count of the mutation itself,
b. The false-positive flip count of its correlated mutations,
c. The false-negative flip number of the mutation itself,
d. The false-negative flip number of its mother mutations in its assigned subclone,
e. The number of mutant cells carrying the mutation withing its subclone. Based on these metrics, mutations were refined as follows:

### Misplaced clusters

If a mutation, along with its correlated mutations, exhibited an excessive number of false-positive flips (a + b > 0.2), the entire group was considered misplaced. In this case, the cluster was pruned and then re-integrated into the phylogeny at a more plausible location using the Bayesian penalty approach mentioned.

### Relocation of orphaned mutations

If a mutation displayed a high false-negative flip count (c > 2) concurrent with a low mutant cell count in its subclone (e < 2), it was deemed to lack a valid phylogenetic parent. Such mutations were relocated to an independent clonal branch.

Mutations that failed to meet the quality thresholds after one iteration of refinement were discarded from the final phylogeny. This process was repeated iteratively 1-5 times until the tree topology stabilized.

### Removal of likely doublet cells

Finally, likely doublet cells contain too many false-positive flips ratio (> 0.2) were excluded from the final tree.

### Benchmarking of PhyloSOLID

For the benchmark, we used real-world single-cell whole-genome sequencing (WGS) data and WGS data from single-cell colonies^18–21^. The ground-truth mutation lists and phylogenies for these datasets were sourced from their original publications. Germline variants and artifact mutations were identified by randomly sampling reads from the raw BAM files using mpileup (Supplementary Table 4). The second simulated dataset used in Fig. 1f was generated by modeling the allele dropout ratios observed in a realistic scRNA-seq data.

For the real-world single-cell RNA-seq data, candidate mosaic mutations were called using MosaicSC^24^, and the ground-truth CRISPR-barcode phylogenies were obtained from the corresponding original papers (Supplementary Table 5-6).

Phylogenetic trees were constructed from both simulated and real-world single-cell data using **eight** software tools, including HUNTRESS, CellPhy, ScisTree, ScisTree2, BSCITE, SiFit, PhISCS and MPBoot, and compared against PhyloSOLID. Default parameters were used for all tools. The tanglegrams and comparison plots between the barcode trees and mutation trees generated by each method were produced using R packages dendextend^33^ (v1.17.1) and phangorn^34^ (v2.11.1).

**Supplementary Fig. 1:**
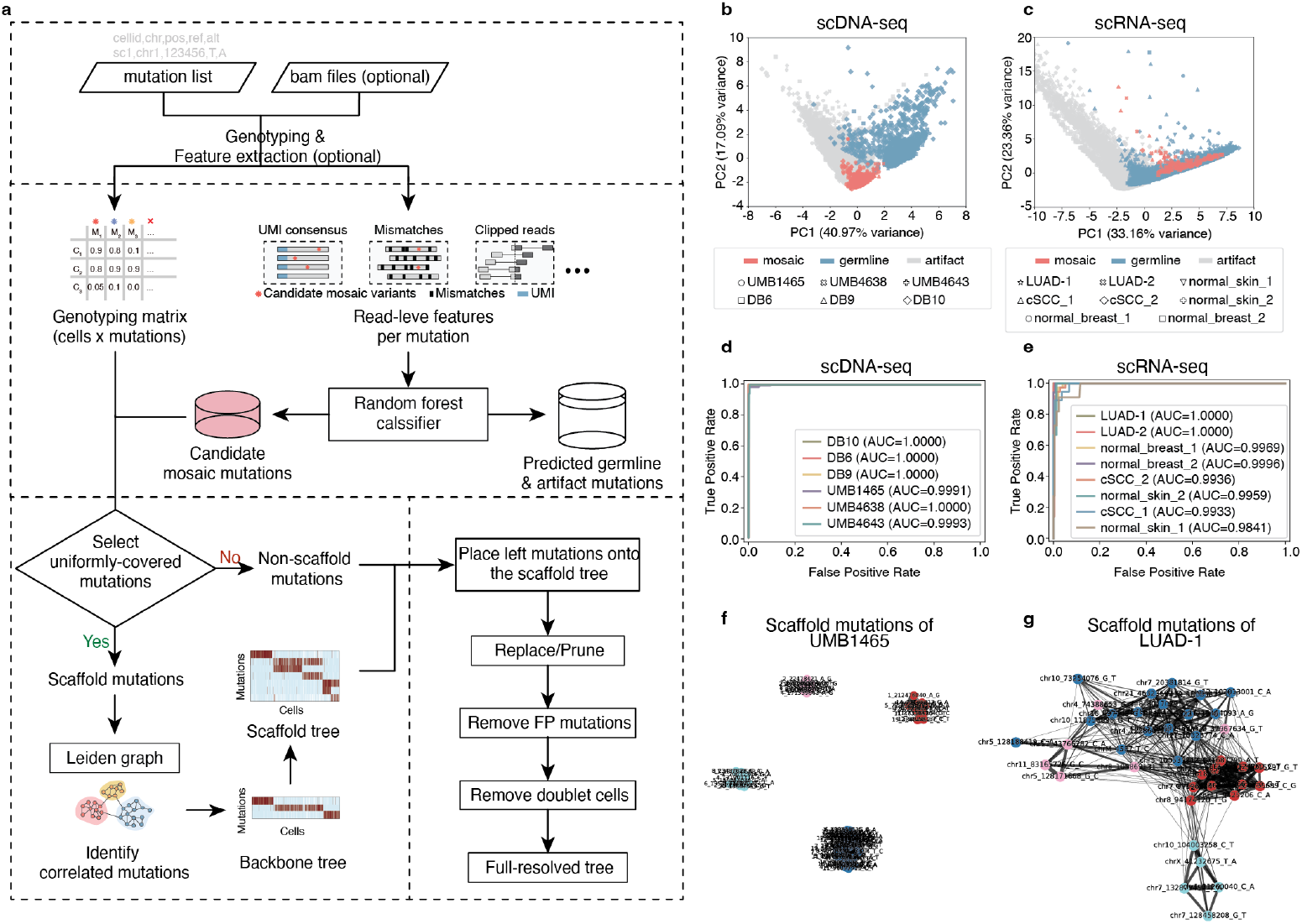
Framework of PhyloSOLID. **a**, Overall pipeline. See Methods for details. **b-c**, Principal component analysis (PCA) of extracted sequence features distinguishes somatic mutations from germline variants and technical artifacts. In the resulting PCA space, true somatic mutations form a distinct cluster, separable from both germline variants and artifacts. **d**-**e**, Application of a logistic regression classifier to discriminate somatic mutations. Receiver operating characteristic (ROC) curves demonstrate high classification accuracy on both scDNA-seq and scRNA-seq benchmark datasets. **f**-**g**, Identification of correlated backbone mutations via a leaden graph. Representative leiden graphs derived from scDNA-seq (**f**) and scRNA-seq (**g**) data are shown. Benchmark dataset details: UMB1465, UMB4638, UMB4643: scDNA-seq data after whole-genome amplification (sample IDs from Lodato et al., *Science*, 2018). DB6, DB9, DB10: Single-cell colony data (sample IDs from Bae et al., *Science*, 2018, and Park et al., *Nature*, 2021, respectively). LUAD-1 and LUAD-2 correspond to the original sample IDs 100k and 10k, respectively (Quinn et al., *Science*, 2021). cSCC_1 (original sample ID: P6_cSCC) and normal_skin_1 (original sample ID: P6_normal) are from Andrew et al., *Cell*, 2020. normal_breast_1 (original sample ID: hbca_n03) and normal_breast_2 (original sample ID: hbca_c11) are from Tapsi et al., *Nature*, 2023.

**Supplementary Fig. 2:**
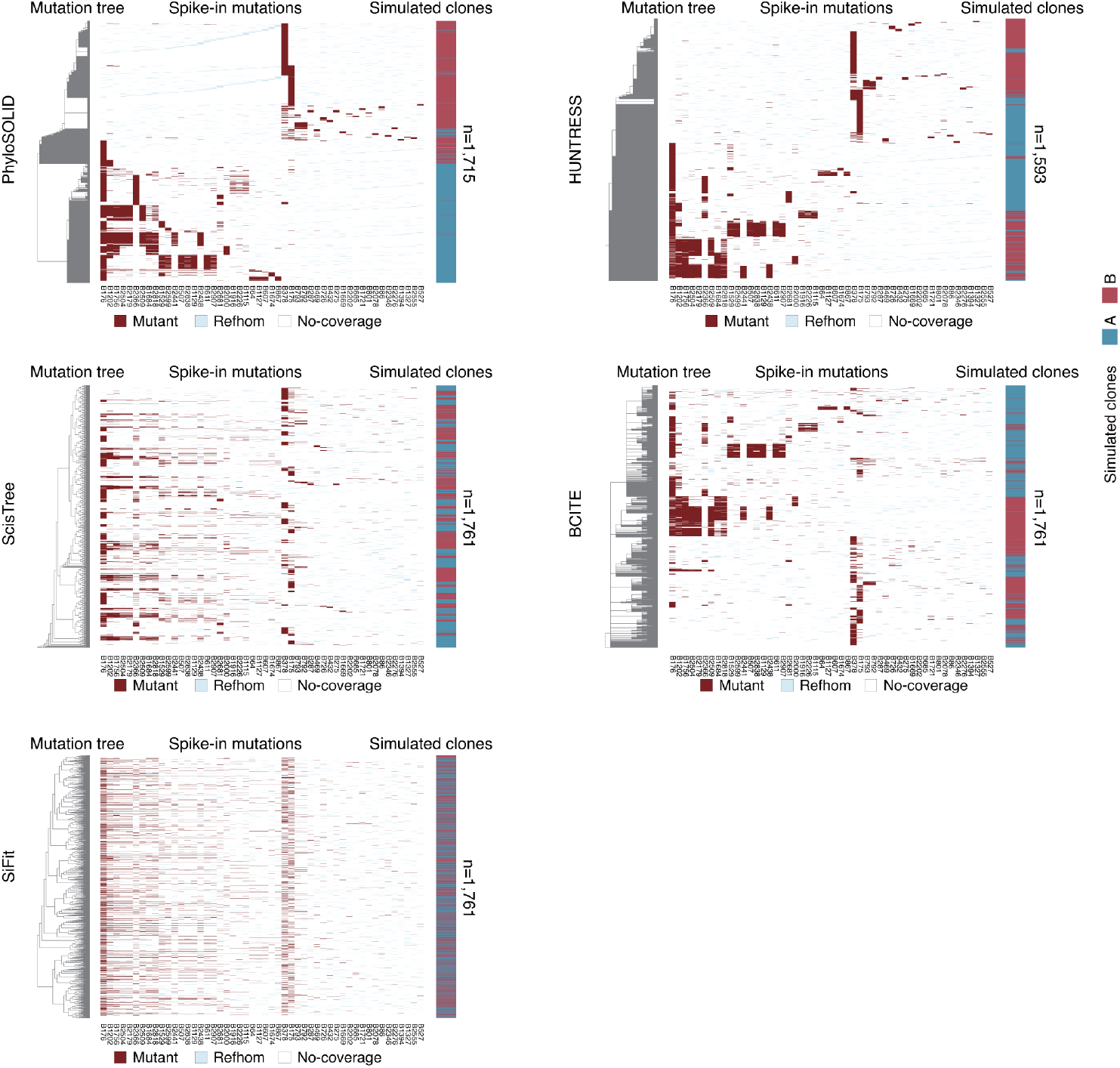
Benchmarking PhyloSOLID on simulated data with high sparsity. The ability of each method to recover the true monoclonal structure under high dropout conditions is assessed. For each panel, phylogeny generated by each software was shown on the left and simulated mutation matrix was shown on the right. Rows correspond to cells and columns to simulated mutations. Mutant cells are shown in red, reference-homozygous (refhom) sites in light blue, and sites with no coverage in white. Execution of PhiSCS, CellPhy, ScisTree2 and MPBoot failed on this dataset.

**Supplementary Fig. 3:**
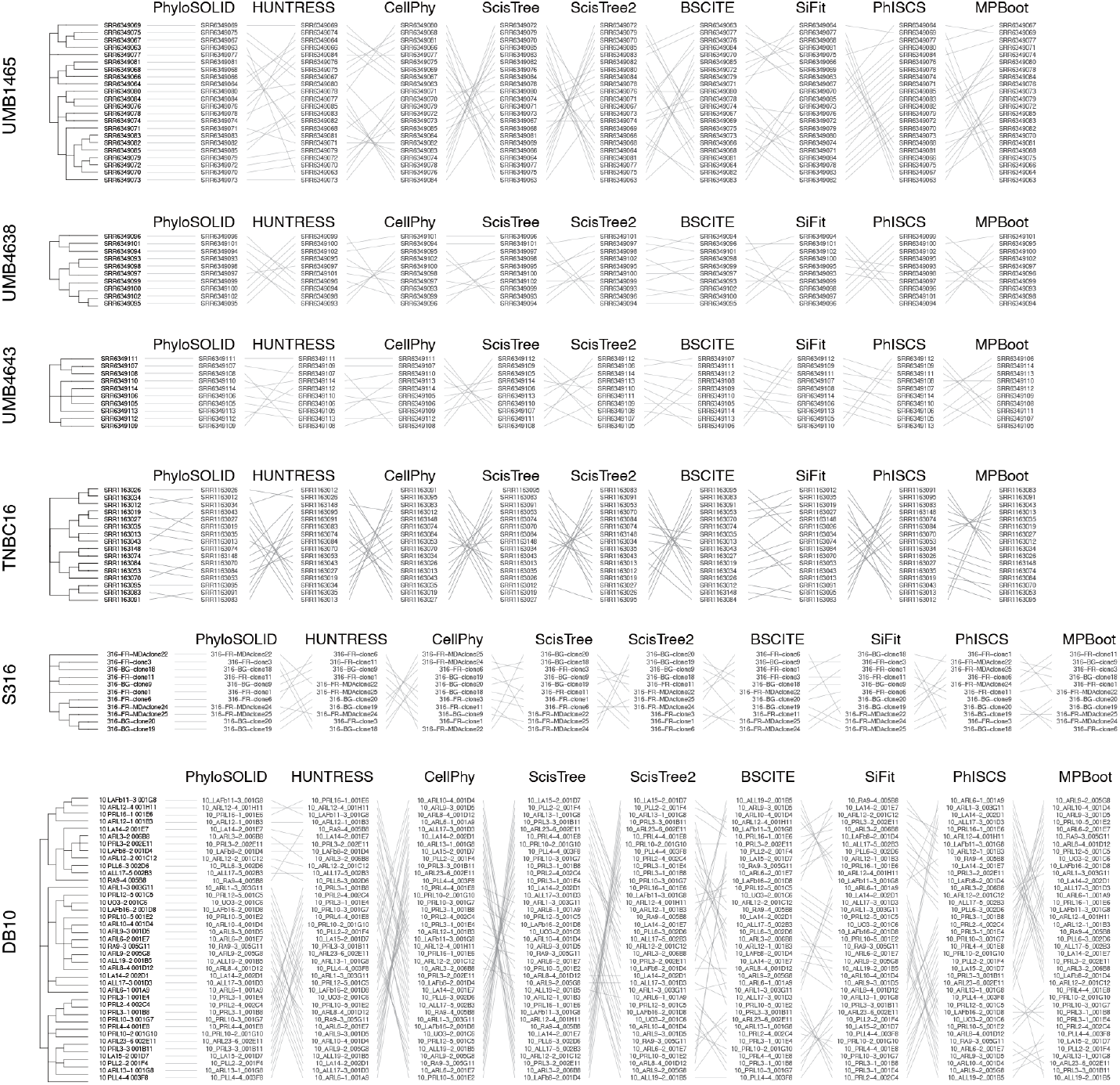
Benchmarking PhyloSOLID on scDNA-seq data. Ground-truth phylogeny (leftmost tree, from published data) and the corresponding phylogenies inferred by each software. Each row represents a cell. Tanglegrams were shown to visualize their agreement with the ground-truth tree (the left-most tree). Benchmark dataset details: UMB1465, UMB4638, UMB4643: scDNA-seq data after whole-genome amplification (sample IDs from Lodato et al., *Science*, 2018). TNBC16: scDNA-seq data after whole-genome amplification (sample ID from Wang et al., *Nature*, 2014). S316, DB10: Single-cell colony data (sample IDs from Bae et al., *Science*, 2018, and Park et al., *Nature*, 2021, respectively).

**Supplementary Fig. 4:**
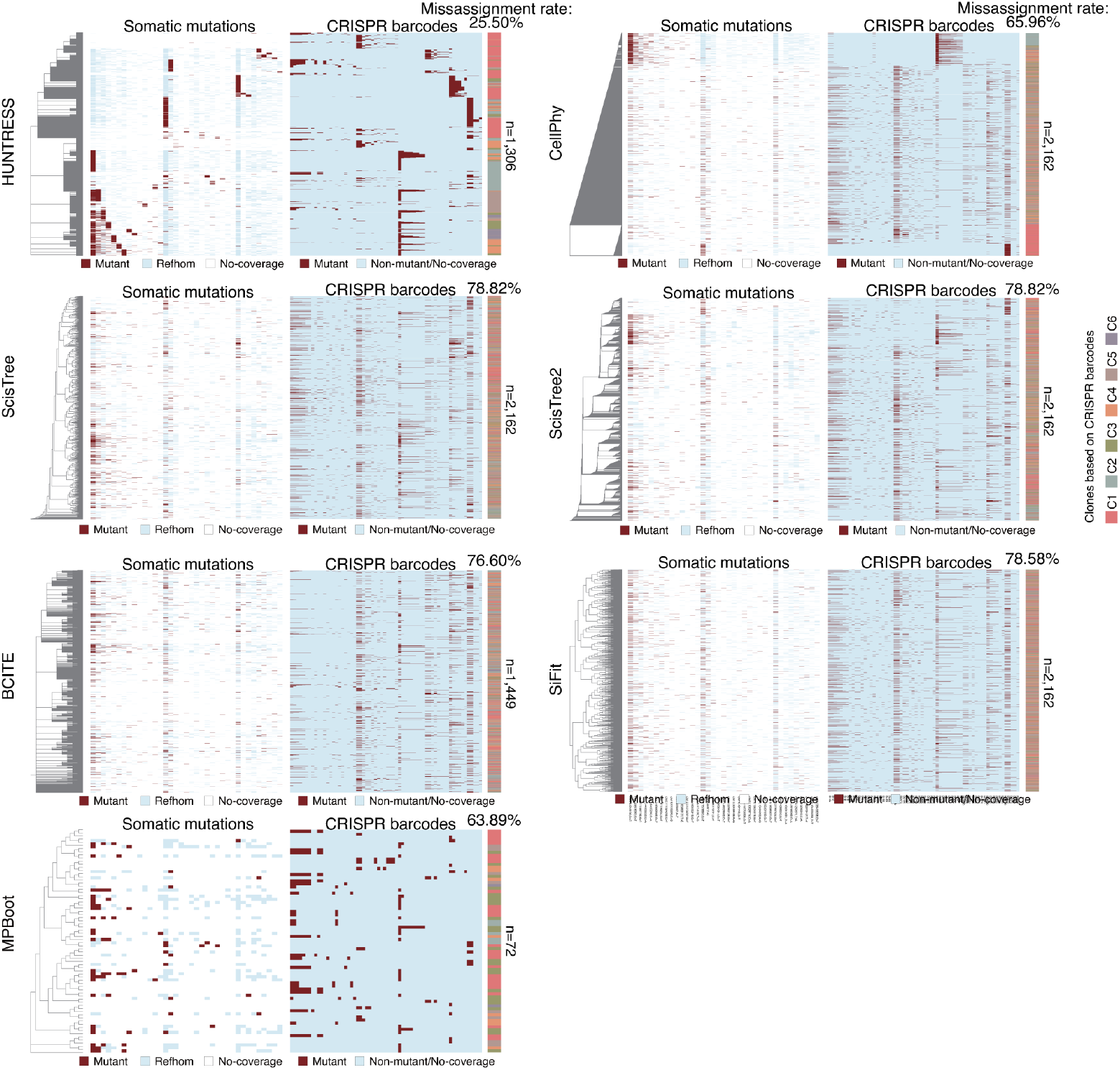
Benchmarking phylogeny reconstruction tools on scRNA-seq data (LUAD-1) with CRISPR barcode ground truth. For each evaluated tool, the output is organized as follows: (Left) Phylogeny inferred from mosaic mutations called from scRNA-seq data. (Middle left) Cell-by-mutation matrix. Rows: cells; columns: mutations. Colors denote mutant cells (red), reference-homozygous sites (light blue), and sites with no coverage (white). (Middle right) Barcode-by-cell matrix. Execution of PhiSCS failed on this dataset. Each column represents one CRISPR barcode mutation. (Right) Published ground-truth clonal structure based on CRISPR barcodes. Benchmark dataset details: the original sample ID of LUAD-1 is “100k” (Quinn et al., *Science*, 2021).

**Supplementary Fig. 5:**
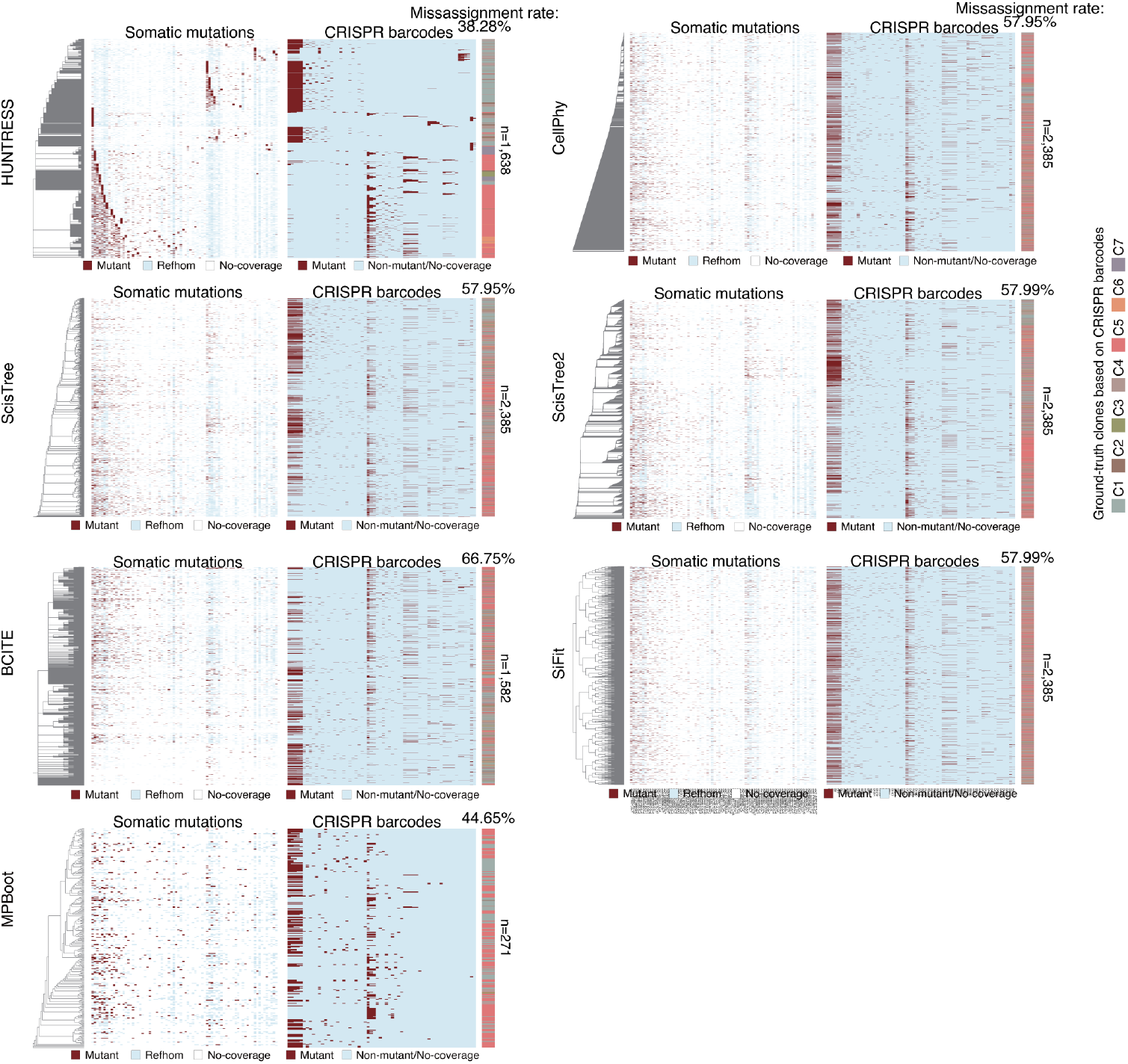
Benchmarking phylogeny reconstruction tools on scRNA-seq data with CRISPR barcode ground truth (LUAD-2). For each evaluated tool, the output is organized as follows: (Left) Phylogeny inferred from mosaic mutations called from scRNA-seq data. (Middle left) Cell-by-mutation matrix. Rows: cells; columns: mutations. Colors denote mutant cells (red), reference-homozygous sites (light blue), and sites with no coverage (white). (Middle right) Barcode-by-cell matrix. Execution of PhiSCS failed on this dataset. Each column represents one CRISPR barcode mutation. (Right) Published ground-truth clonal structure based on CRISPR barcodes. Benchmark dataset details: the original sample ID of LUAD-2 is “10k” (Quinn et al., *Science*, 2021).

**Supplementary Fig. 6:**
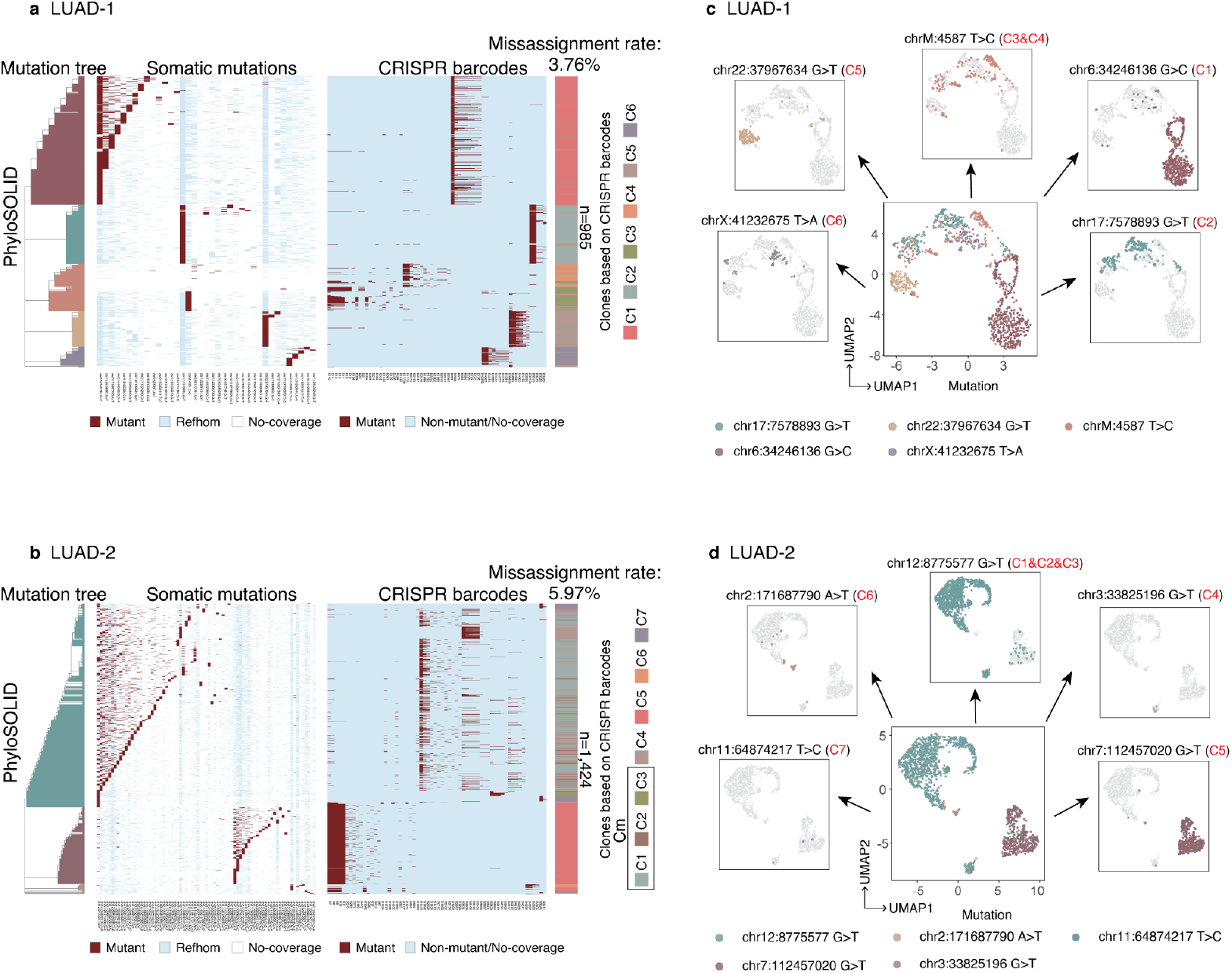
Benchmarking PhyloSOLID on scRNA-seq data with CRISPR barcode ground truth. **a, b**, Analysis of two lung cancer scRNA-seq datasets. For each dataset: (Left) Phylogeny reconstructed from mosaic mutations identified in the scRNA-seq data. (Middle left) Cell-by-mutation matrix. Rows represent cells, columns represent mutations. Mutant cells are shown in red, reference-homozygous (refhom) sites in light blue, and sites with no coverage in white. (Middle right) Barcode-by-cell matrix. Each column corresponds to one CRISPR barcode mutation. (Right) Ground-truth clonal structure derived from the published CRISPR barcode data. **c, d**, UMAP visualization of cells colored by their assigned clones. Benchmark dataset details: the original sample IDs of LUAD-1 and LUAD-2 are “100k” and “10k” (Quinn et al., *Science*, 2021).

